# LSD reconfigures the frequency-specific network landscape of the human brain

**DOI:** 10.1101/2025.03.21.644645

**Authors:** Kenneth Shinozuka, Mattia Rosso, Clayton R. Coleman, Leor Roseman, Mendel Kaelen, Suresh Muthukumaraswamy, David J. Nutt, Robin Carhart-Harris, Peter Vuust, Morten L. Kringelbach, Leonardo Bonetti

## Abstract

Lysergic acid diethylamide (LSD) and other psychedelic substances profoundly alter human consciousness. While several studies have demonstrated changes in brain function and connectivity associated with psychedelics, we still have a limited understanding of how LSD reshapes brain networks operating across different frequency bands. In this study, we applied the recently developed FREQ-NESS method to MEG data from 14 healthy participants who received LSD under four conditions: eyes-closed with or without music and eyes-open with or without a video stimulus. LSD significantly restructures canonical networks in the alpha and beta bands. Relative to broadband brain activity, it enhances the prominence of high alpha (12.1, 13.3 Hz) across all experimental conditions and high beta (25.3 Hz) in three conditions. Conversely, LSD decreases the prominence of low beta (18.1, 19.3 Hz) in both Open and Closed conditions and low alpha (8.5 Hz) in the latter. In addition, LSD substantially alters the spatial distributions or topographies of the networks. Under LSD, the low alpha (8.5 Hz) network shifts anteriorly toward the motor cortex, while high alpha (12.1, 13.3 Hz) becomes more localized to the visual cortex. Low beta (18.1, 19.3 Hz) expands over the temporal and occipital cortices, whereas high beta (25.3, 26.5 Hz) topographies remain unchanged. Our findings provide critical insights into the specific frequencies and spatial networks in which LSD modulates brain connectivity, adding nuance to prevailing theories about network disintegration under psychedelics.

**Significance statement:** Psychedelic substances such as lysergic acid diethylamide (LSD) profoundly alter perception and cognition, yet their effect on brain activity across different frequency bands remains unclear. Using FREQ-NESS applied to MEG data, we show that LSD reorganizes brain networks by enhancing high-frequency alpha and beta rhythms while suppressing lower-frequency counterparts. These changes occur across various experimental conditions and shift the spatial distribution of brain activity, particularly in motor and visual regions. Our findings suggest that LSD modulates brain connectivity in a frequency- and region-specific manner, offering new insights into how psychedelics alter consciousness. This work advances our understanding of LSD’s neural effects, which may be relevant for therapeutic applications and models of brain function.

## Section 1. Introduction

Psychedelic drugs, including lysergic acid diethylamide (LSD), psilocybin (the main psychoactive ingredient in magic mushrooms) and N,N-dimethyltryptamine (DMT), have recently been reappraised as potentially effective treatments for a range of psychiatric conditions, including generalized anxiety disorder and depression (Beutler et al., 2024; D’Souza et al., 2022; Goodwin et al., 2022; Holze et al., 2023; Raison et al., 2023; Shinozuka, Tabaac, et al., 2024; Tabaac et al., 2024). All “classic” psychedelics are agonists at the serotonin-2A or 5-HT_2A_ receptor, and blocking this receptor subdues or eliminates their hallucinogenic effects (Preller et al., 2017; Quednow et al., 2012; Vollenweider et al., 1998). While the neural mechanisms of psychedelics are not fully understood, one consistent result in the functional neuroimaging literature is that these drugs “disintegrate and desegregate” brain networks (Carhart-Harris et al., 2016; Dai et al., 2023; Madsen et al., 2021; Mason et al., 2020; Müller et al., 2018; Shinozuka, Jerotic, et al., 2024; Timmermann et al., 2023). That is, in functional magnetic resonance imaging (fMRI) studies, psychedelics increase functional connectivity between networks while decreasing it within networks.

However, fMRI is limited by its low temporal resolution. Faster modalities like electroencephalography (EEG) and magnetoencephalography (MEG) have demonstrated not only that brain networks arise and disperse within a span of 100-200 milliseconds, but also that different networks operate on different timescales (Baker et al., 2014; Gohil et al., 2022). For instance, the visual network oscillates primarily at alpha frequencies, whereas the anterior default mode network does so at delta frequencies (Gohil et al., 2022). Additionally, MEG is better suited for measuring the neural correlates of psychedelics than fMRI, since the vasoconstrictive effects of psychedelics confound the fMRI but not the MEG signal (Dyer & Gant, 1973; Hillman, 2014; Özbay et al., 2019; Wilson et al., 2016).

While many EEG/MEG studies have found that psychedelics reduce power across multiple frequency bands (Carhart-Harris et al., 2016; Eidelberg et al., 1965; Horovitz et al., 1965; Hutten et al., 2024; Le et al., 2024; Murray et al., 2022; Muthukumaraswamy et al., 2013; Rodin & Luby, 1966; Vejmola et al., 2021), very few studies to date have investigated the effects of psychedelics on brain *networks* within specific frequency bands. Muthukumaraswamy et al. (2013) used independent component analysis (ICA) to extract frequency-resolved brain networks from MEG data on psilocybin. They found that psilocybin reduced the activation of seven different networks, including bilateral frontoparietal, motor, and visual networks, in a range of frequency bands.

However, a problem with traditional power and ICA analyses is that they do not inherently correct for broadband activity (Gyurkovics et al., 2021). Due to this confounding effect, it is unclear whether the results of these analyses can be attributed to changes in power or connectivity within specific frequency bands, or merely to “background” broadband noise. These analyses are therefore especially unreliable for interventions like psychedelics, which affect power in a broadband manner. Additionally, ICA does not provide information on the variance associated with different components, making it challenging to determine how many components and which ones are of interest. Furthermore, the MEG-ICA study on psychedelics focused on canonical frequency bands, neglecting a proper frequency-resolved approach, which is crucial for detecting subtle yet significant changes in brain connectivity (Rosso et al., 2025).

A new framework, the FREQuency-resolved variant of Network Estimation via Source Separation (FREQ-NESS), offers a solution to these problems by isolating frequency-resolved from broadband connectivity in source space reconstructed with an 8 mm grid (Rosso et al., 2025). It does so by determining networks that maximally separate the broadband covariance from the narrowband covariance, i.e., the covariance of MEG data that has been bandpass filtered to individual frequencies. Prior applications of FREQ-NESS recapitulated canonical resting-state networks at specific frequencies, including occipital alpha and sensorimotor beta networks (Rosso et al., 2025). Intriguingly, playing a 2.4 Hz sound significantly increased the prominence of brain activity at 2.4 Hz and shifted the spatial configuration of the corresponding networks towards auditory cortex.

Here, we apply FREQ-NESS to previously acquired MEG data on the acute effects of LSD (Carhart-Harris et al., 2016). Fourteen healthy human participants were administered both LSD and placebo in two separate sessions. Resting-state data were recorded across four experimental conditions: (1) eyes-open resting-state, (2) eyes-closed resting-state, (3) listening to ambient music with eyes closed, (4) watching a silent video. Our aim is to determine the frequency-specific brain networks that characterize the LSD state, as well as the changes in the prominence of different frequencies. We also aim to understand how the reorganization of brain networks is modulated by auditory and visual stimulation, and their interaction with the effect of LSD. These networks may provide novel, comprehensive perspectives on the neural mechanisms of the hallucinations and altered states of consciousness that psychedelics elicit.

## Section 2. Methods

### Section 2.1. Data acquisition

The data was acquired in a previous study (Carhart-Harris et al., 2016), and the experimental protocol is described in detail in the supplementary materials therein. This study was approved by the National Research Ethics Service committee London-West London and was conducted in accordance with the revised declaration of Helsinki (2000), the International Committee on Harmonization Good Clinical Practice guidelines, and National Health Service Research Governance Framework.

Each participant underwent two scanning sessions, one in which they received placebo and another in which they received 75 μg of intravenously administered LSD. Approximately 165 minutes after the LSD was administered, the participants were recorded with a CTF 275-gradiometer MEG at a sampling rate of 600 Hz, though four of the sensors were turned off because of excessive sensor noise. (fMRI data was also collected, but it was not analysed in this study.) During the MEG recording, participants were subjected to four conditions: eyes-open resting-state (referred to in the paper as “Open”), eyes-closed resting-state (“Closed”), listening to music with eyes closed (“Music”), and watching a silent nature documentary (“Video”). Each scan lasted approximately seven minutes. While there was one scan associated with each of the Music and Video conditions, two scans were associated with each of the Open and Closed conditions; therefore, we randomly selected one Open and one Closed scan for a total of four scans per subject.

Of the original 20 participants, five were excluded from the data analysis because they either failed to complete both scanning sessions (placebo and LSD) or because their movement artefacts were too large. Additionally, one participant failed to complete all four conditions; they were missing data from the Music condition. Therefore, data from 14 participants were analysed in this study.

### Section 2.2. Data preprocessing and source reconstruction

Data was high-pass filtered to 1 Hz and then downsampled to 200 Hz. The logistic infomax algorithm of independent component analysis (ICA) was used to identify eye blink and heartbeat artefacts (Bell & Sejnowski, 1995). Components containing these artefacts were manually removed by visual inspection. Note that the root mean square of the electrocardiography (ECG) and electrooculography (EOG) signals did not significantly differ between placebo and LSD, suggesting that the drug did not affect the magnitude of these artefacts. There was also no significant difference between placebo and LSD in the number of ICA components that were removed. Note that the data was preprocessed differently than in the original study, in which artifacts in the MEG data were also removed with other manual and automatic methods. For the sake of this manuscript, we deemed that additional artifact removal was either unnecessary or redundant with ICA. Additionally, as mentioned above, participants with strong movement artifacts were already excluded prior to preprocessing, based on previous analyses.

To perform source reconstruction, an 8 mm^3^ grid containing 3559 voxels in MNI space was inversely warped to each subject’s native-space anatomical MRI. A linearly constrained minimum variance (LCMV) beamformer was applied to the inversely-warped template, with the regularization parameter set to 5% of the average of the diagonal elements of the sensor covariance matrix (Van Veen et al., 1997). For the dipole at each centroid in the source model, only the orientation that maximizes power was used to estimate the spatial filter of the beamformer.

### Section 2.3. FREQ-NESS

FREQ-NESS uses generalized eigendecomposition (GED) to identify brain networks that are associated with particular frequencies (Cohen, 2017, 2022; Rosso et al., 2021, 2022, 2023). In particular, for each frequency of interest, GED determines the voxel weights that maximally separate the covariance of the signal narrow band filtered to that frequency from the covariance of the broadband signal. This technique therefore decomposes the networks that are *specific* to individual frequencies. This is accomplished by solving the following equation for each frequency of interest:

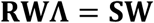

where **W** is a 3559 x 3559 matrix of eigenvectors containing the aforementioned voxel weights, **∧** is a 3559 x 3559 diagonal matrix containing the associated eigenvalues in the main diagonal, **R** is the broadband covariance matrix (3559 x 3559), and **S** is the narrow-band covariance matrix at the frequency of interest (3559 x 3559).

When normalized in percent units, each eigenvalue can be interpreted naturally as the proportion of variance explained by the corresponding eigenvector. The eigenvectors are then projected into voxel space. That is, for each such eigenvector **w**, the corresponding network activation pattern **a** is:

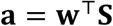

This step effectively reweights the eigenvectors so that they are physiologically interpretable as the spatial distribution of network activation across voxels (Haufe et al., 2013).

Narrow-filtering the MEG signal is accomplished by applying a Gaussian wavelet kernel with a width of 0.3 Hz at half of the maximum gain, centered at each frequency of interest. The frequencies of interest were a sample of 86 frequencies ranging from 0.2 to 97.6 Hz, generally spaced out by 1.2 Hz. The width of the filter was optimized for every frequency based on a log10 scale. For further details on the derivation and implementation of FREQ-NESS, the reader is referred to the reference text (Rosso et al., 2025).

### Section 2.4. Statistical tests

A three-way repeated-measures Analysis of Variance (ANOVA) was used to measure statistically significant changes in the leading eigenvalue. Results from only 14 participants per condition were inputted into the ANOVA, since data from only 14 participants was included in our analysis of the Music condition. Within-subject factors were Frequency, Condition, and Drug. If applicable, post-hoc *t*-tests with Bonferroni correction were conducted to determine pairwise, between-drug differences in the leading eigenvalue at each frequency and condition. If the assumption of sphericity was violated, Greenhouse-Geisser corrections were automatically applied to adjust the degrees of freedom. The rstatix package in R was used to conduct all statistical analyses.

### Section 2.5. Correlations with subjective ratings

For each frequency that exhibited a significant between-drug difference in the leading eigenvalue, this difference was correlated with responses to questions about six subjective experiences. These questions were: 1) “Please rate the intensity of the drug effects during the last scan”, with a bottom anchor of “no effects”, a mid-point anchor of “moderately intense effects” and a top anchor of “extremely intense effects”; 2) “With eyes closed, I saw patterns and colours”, with a bottom anchor of “no more than usual” and a top anchor of “much more than usual”; 3) “With eyes closed, I saw complex visual imagery”, with the same anchors as item 2; 4) “How positive was your mood for the last scan?”, with the same anchors as item 2, plus a mid-point anchor of “somewhat more than usual”; 5) “I experienced a dissolving of my self or ego”, with the same anchors as item 2; and 6) “Please rate your general level of emotional arousal for the last scan”, with a bottom anchor of “not at all emotionally aroused”, a mid-point anchor of “moderately emotionally aroused” and a top anchor of “extremely emotionally aroused”. Ratings of these six items were obtained immediately after each MEG scan on a visual analog scale (VAS), with a minimum rating of 0 and a maximum of 20. Using False Discovery Rate (FDR), Spearman correlations between the leading eigenvalue and VAS ratings were corrected for multiple comparisons across questionnaire items, experimental conditions, and frequencies of interest.

## Section 3. Results

### Section 3.1. LSD alters the prominence of alpha and beta networks

The FREQ-NESS method uses generalized eigendecomposition to decompose MEG voxel timeseries into brain networks that are associated with specific frequencies. At each frequency, the leading eigenvector, i.e., the eigenvector associated with the largest eigenvalue, contains the voxel weights of the brain network that explains the most variance relative to the broadband covariance. We define the “leading component” as the network activation pattern associated with the leading eigenvector. The leading eigenvalue represents the amount of variance explained by the leading component.

Our three-way repeated-measures ANOVA identified a significant main effect of frequency (*F*_29,377_ = 182.885, *p* < 0.0001) and condition (*F*_1.77,23.04_ = 9.090, *p* = 0.002) on the leading eigenvalue, as well as a significant interaction effect of frequency by condition (*F*_87,1131_ = 46.312, *p* < 0.0001) and of frequency by drug (*F*_29,377_ = 2.604, *p* < 0.0001). (The main effect of drug tended towards significance [*F*_1,13_ = 3.641, *p* = 0.08].) Post-hoc multiple comparisons tests revealed that, in all four conditions, LSD significantly increases the amount of variance explained by the leading component at 12.1 Hz (high alpha; *p*_Open_ = 0.010, *p*_Closed_ = 0.0005, *p*_Music_ = 0.009, *p*_Video_ = 0.032) and at 13.3 Hz (high alpha; *p*_Open_ = 0.009, *p*_Closed_ = 0.003, *p*_Music_ = 0.016, *p*_Video_ = 0.003) (**Figure 2**). At 25.3 Hz (high beta), the leading eigenvalue also increased significantly on LSD in three of the conditions (*p*_Open_ = 0.049, *p*_Music_ = 0.044, *p*_Video_ = 0.031). LSD significantly decreased the leading eigenvalue at 18.1 Hz (low beta) in two conditions (*p*_Open_ = 0.014, *p*_Closed_ = 0.01), as well as at 8.5 Hz (low alpha) and 19.3 Hz (low beta) in just the Closed condition (*p*_Closed_ = 0.034 and *p*_Closed_ = 0.003, respectively). Finally, the leading eigenvalue significantly increased on LSD at 26.5 Hz (high beta) in the Video condition (*p*_Video_ = 0.02).

**Figure 1.**
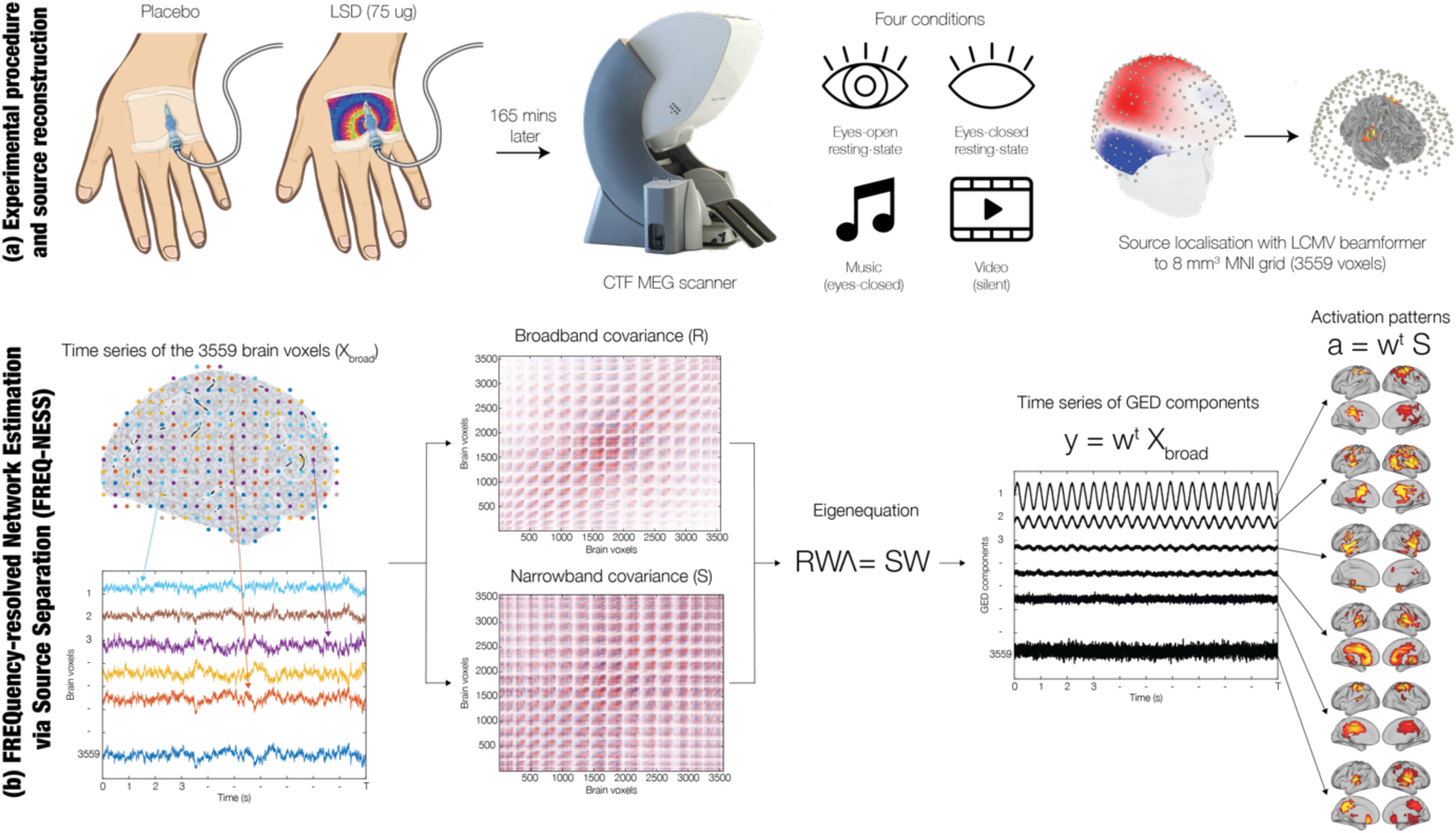
Overview. (a) Fourteen human participants were each administered placebo and 75 micrograms (ug) of LSD in two separate sessions (Carhart-Harris et al., 2016). Each session included four separate conditions: eyes-open resting-state (hereafter referred to as “Open”), eyes-closed resting-state (“Closed”), listening to music with eyes closed (“Music”), and watching a silent video (“Video”). After a minimal preprocessing pipeline that included high-pass filtering, downsampling, and artefact removal with ICA, the MEG channel timeseries were source reconstructed to an 8 mm^3^ grid in MNI space with a linearly-constrained minimum variance (LCMV) beamformer. (b) The source reconstruction yields 3559 brain voxel timeseries. These timeseries are narrow-band filtered at a range of 86 equally spaced frequencies. For each frequency, the narrowband covariance matrix, S, is computed. The sets of voxel weights W that maximally separate S from the broadband covariance matrix, R, are determined by solving the eigenequation RWΛ = SW, where Λ contains the variance explained by each set of voxel weights. W is projected into voxel space to obtain the network activation patterns, or spatial topographies, A, and the network timeseries can also be computed. We refer to the network that explains the most variance at a given frequency as the “leading component.”

**Figure 2.**
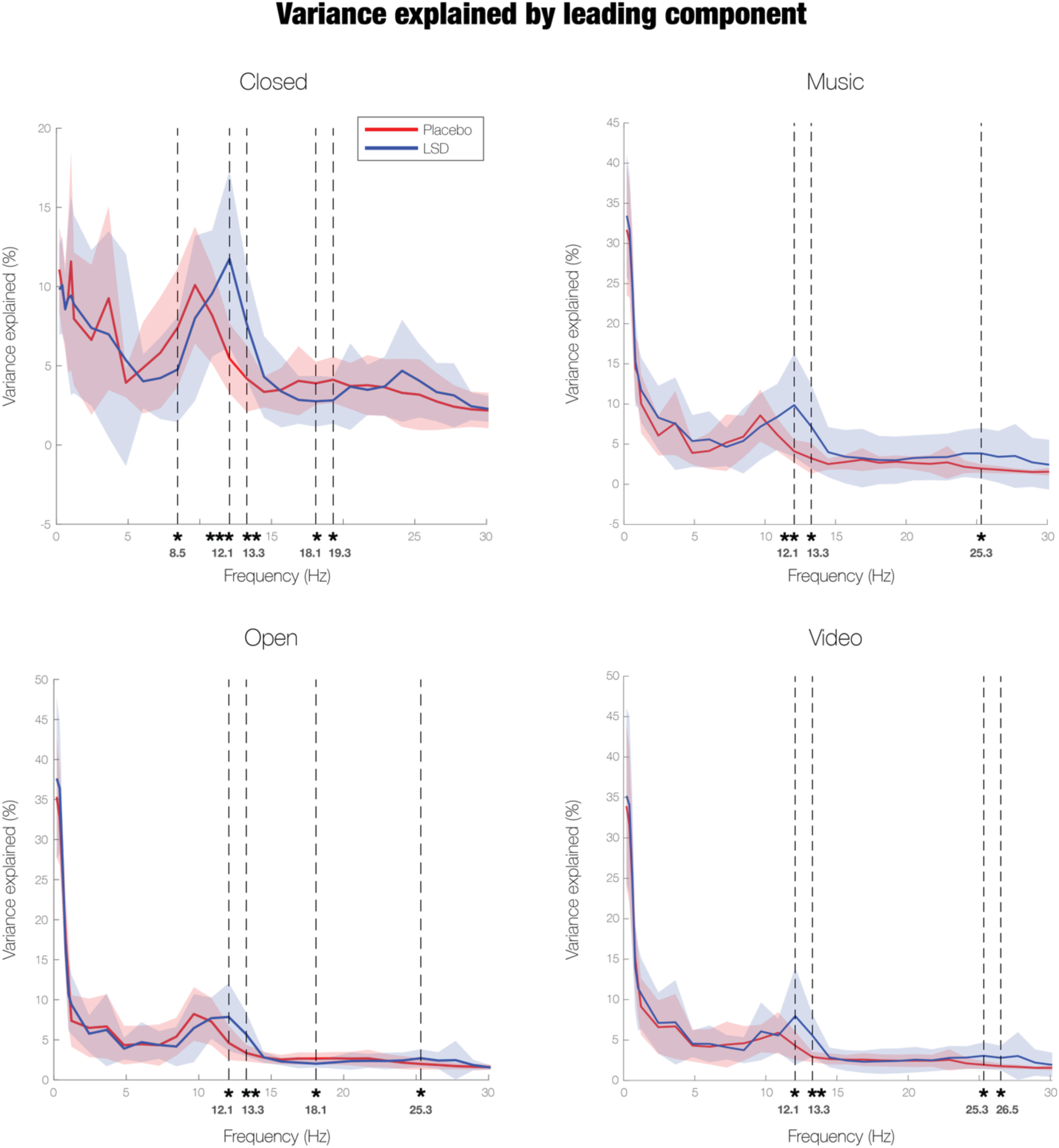
LSD significantly increases the amount of variance explained by high alpha activity in all four conditions. At 12.1 Hz and 13.3 Hz, the amount of (co)variance captured by the leading component, relative to broadband covariance, grows significantly larger on LSD in the Open, Closed, Music, and Video condition. The same is true of high-beta activity (25.3 Hz) in three of the conditions: Music, Open, and Video. However, in the Open and Closed conditions, low-to mid-beta activity (18.1 Hz) accounts for significantly less of the covariance in the MEG data on LSD. The amount of variance explained by alpha activity (8.5 Hz) also decreases significantly in the Closed condition.

None of the correlations between the subjective ratings and the changes in leading eigenvalue survived multiple comparisons, though several correlations were significant at α = 0.05 before FDR correction. These included correlations between the leading eigenvalue at 25.3 Hz and emotional arousal in the Open condition; and correlations between the leading eigenvalue at 13.3 Hz and emotional arousal, between 19.3 Hz and complex visual hallucinations, and between 19.3 Hz and ego dissolution in the Closed condition.

### Section 3.2. LSD shifts the spatial topographies of frequency-specific brain networks

We now discuss the spatial topographies of the networks at each frequency and condition for which there is a significant between-drug difference in the leading eigenvalue. At low alpha (8.5 Hz), the leading component in the Closed condition expands into more anterior regions on LSD, namely the right primary motor cortex, right primary somatosensory cortex, right supplementary motor area, bilateral middle cingulate, and various bilateral temporal cortices (**Figure S1**). The network withdraws from visual areas such as bilateral V1, bilateral V2, and right superior and middle occipital gyrus.

However, LSD causes the network at a high alpha frequency (12.1 Hz) to move in the opposite direction; that is, the leading components in both eyes-closed conditions (Closed and Music) become more localized to the occipital cortex (**Figure 3**). In both conditions, the network becomes more activated in V1, bilateral V2, and the right middle and superior occipital gyrus. The network also shifts away from the left primary motor and somatosensory cortex and left superior temporal gyrus in both conditions, as well as the left supplementary motor area in the Closed condition and the left middle cingulate in the Music condition. The leading component of the 12.1 Hz network in the Open condition follows a similar pattern on LSD, engaging some left-hemisphere visual areas while disengaging the left primary motor cortex, bilateral primary somatosensory cortex, and bilateral temporal cortices. On the other hand, in the Video condition, the opposite topography emerges; the 12.1 Hz network extends into bilateral primary motor and somatosensory cortex and bilateral supplementary motor area, as well as some frontal regions, while moving away from regions in several levels of the visual cortex hierarchy and temporal cortices.

**Figure 3.**
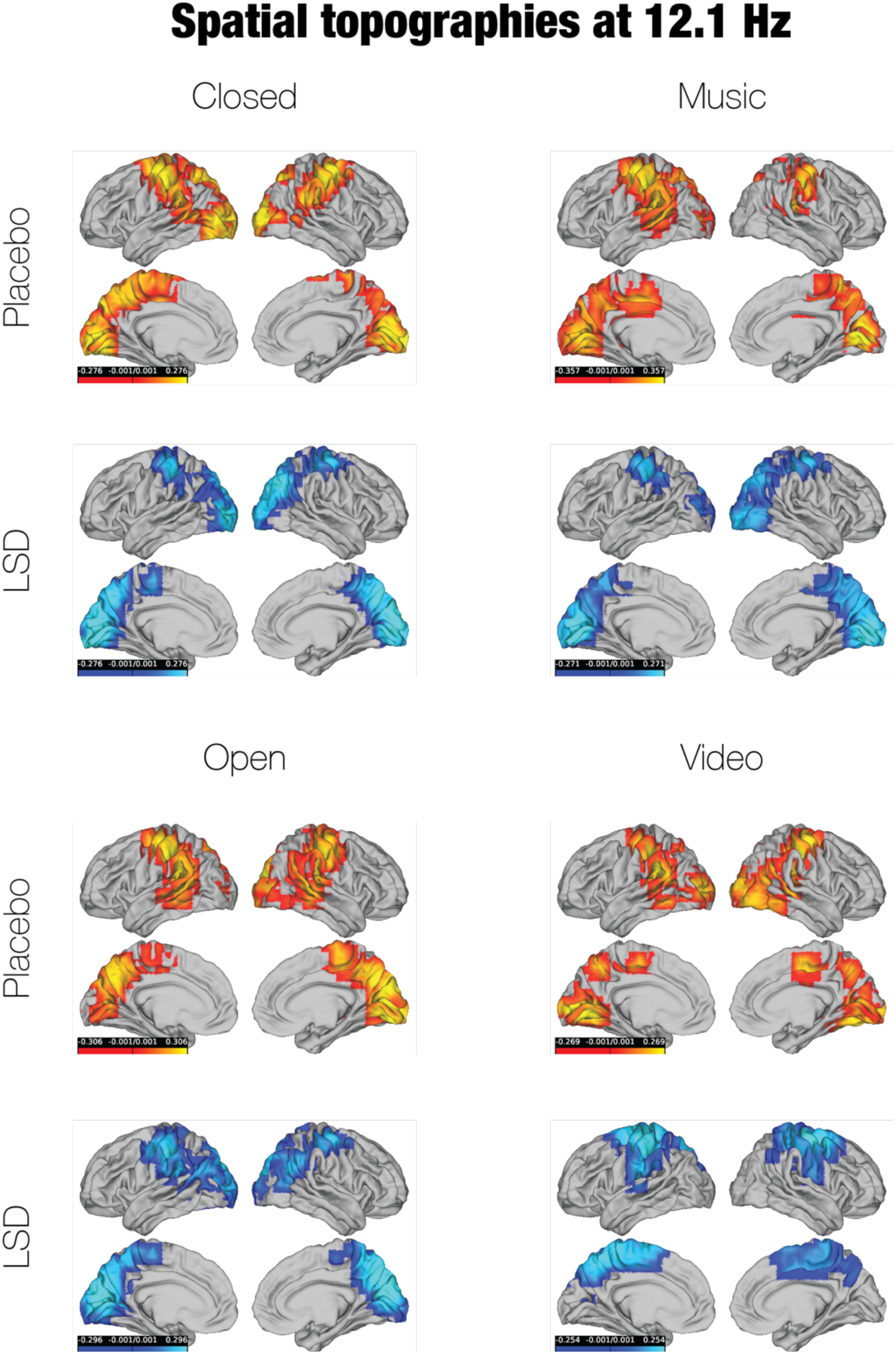
LSD causes the brain network associated with 12.1 Hz activity to become more localised to the occipital cortex in the eyes-closed conditions (Closed and Music), yet more localised to the motor cortex in the Video condition. In the Closed and Music conditions, the spatial topography of the leading component at 12.1 Hz shifts towards visual areas (specifically, V1, V2, and the right middle and superior occipital gyrus) and away from motor areas (specifically, left primary motor and somatosensory cortex). Some, but not all, of these visual areas are also engaged in the 12.1 Hz network in the Open condition, while the same motor areas are disengaged. However, the 12.1 Hz network in the Video condition exhibits the opposite topography; it becomes more concentrated in the bilateral primary motor and somatosensory cortex and less concentrated in bilateral visual areas.

This posterior-to-anterior shift also occurs in a network at the edge of alpha and beta (13.3 Hz) in the Video condition; on LSD, the network recruits the bilateral primary motor and somatosensory cortex, left paracentral lobule (the part of the primary motor cortex that regulates the lower limbs of the body), and various frontoparietal areas, while withdrawing from nearly all visual regions and the temporal lobe (**Figure 4**). While the 13.3 Hz networks in the other conditions also expand into motor cortex and other frontoparietal areas, they nevertheless continue to activate the occipital cortex, as they did at 12.1 Hz. Both the Open and Closed conditions engage right V2 and right middle and superior occipital gyri more on LSD, while the 13.3 Hz network of the Music condition spreads into left V2. On placebo, the 13.3 Hz networks in all four conditions encompass more of the temporal lobe than the 12.1 Hz networks did. However, the 13.3 Hz networks on LSD do not tend to comprise temporal regions.

**Figure 4.**
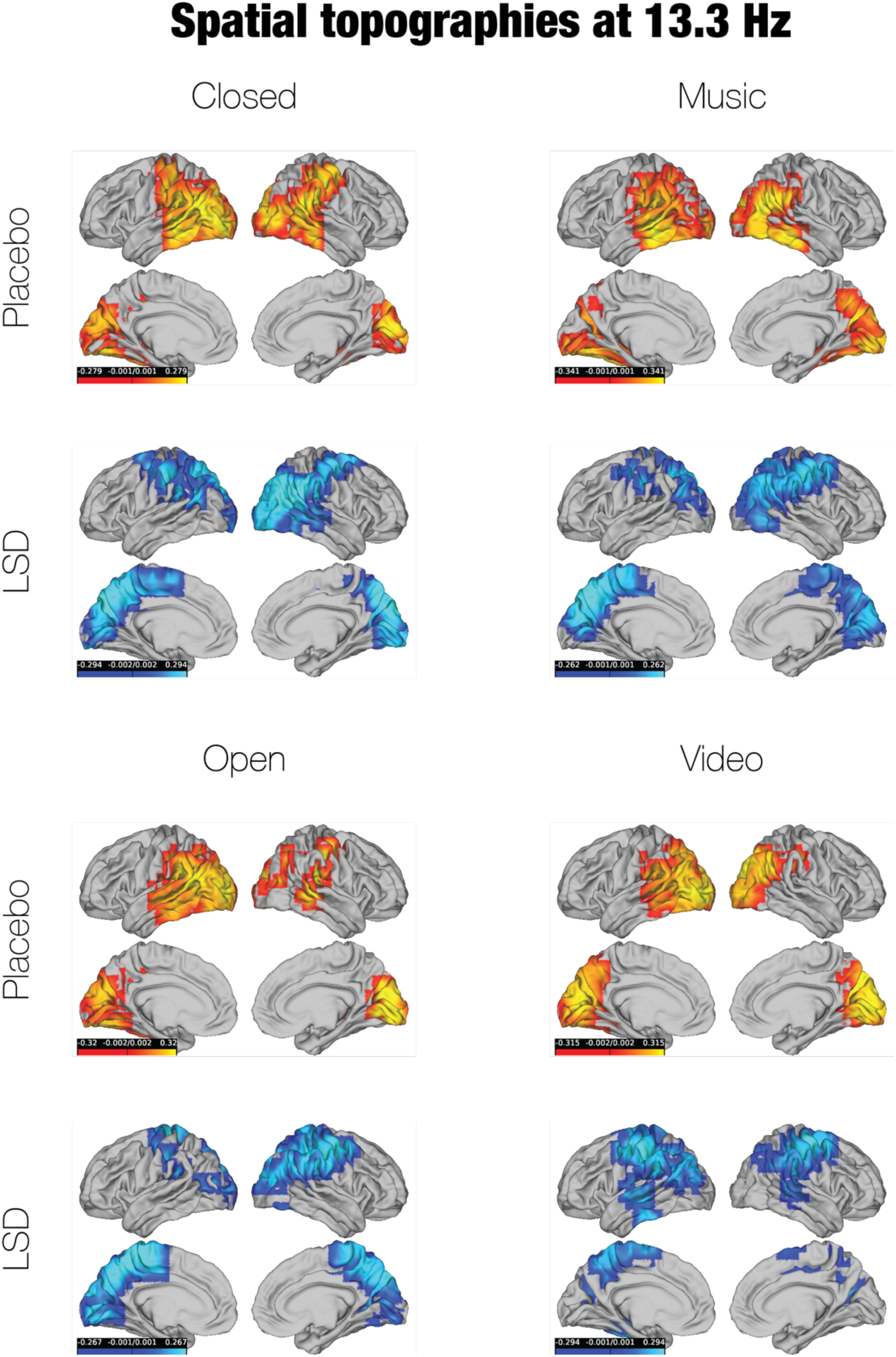
On LSD, the networks associated with 13.3 Hz activity remain active in the occipital cortex, except in the Video condition. In the Open, Closed, and Music conditions, LSD maintains or elevates the activation of some visual areas in the 13.3 Hz network, while also recruiting motor and other frontoparietal areas. However, in the Video condition, the leading component spreads into the bilateral primary motor and somatosensory cortex on LSD and moves away from regions at all levels of the visual cortex hierarchy.

For low beta frequencies, the most consistent pattern is that LSD shifts the leading component towards both left temporal and occipital cortices. LSD shifts the 18.1 Hz network of both the Closed and the Open conditions into the left superior and middle and right middle temporal gyri, as well as the left middle occipital gyrus (**Figure S2**). The 18.1 Hz network of the Closed condition also recruits left V2, left superior occipital gyrus, and right middle occipital gyrus. Similarly, at 19.3 Hz, LSD engages the left superior and middle temporal gyri and the left superior and middle occipital gyri, though the network also moves towards more frontal areas and withdraws from right occipital cortex (**Figure S3**).

Finally, the high beta (25.3 Hz) networks do not exhibit substantial changes between placebo and LSD in the Open, Music, or Video conditions (**Figure S4**). Under both placebo and LSD, the networks are primarily activated in the bilateral primary motor and somatosensory cortex, bilateral supplementary motor area, and other frontal regions, across the three conditions. LSD does not have much of an effect either on the 26.5 Hz network in the Video condition, though the network is less activated in several frontal areas compared to placebo (**Figure S5**).

## Section 4. Discussion

### Section 4.1. Prominence of brain networks on LSD is distinct from power

In this study, we applied FREQ-NESS, an analysis pipeline based on generalized eigendecomposition, to determine the effect of LSD on frequency-specific brain networks. Our main finding is that LSD restructures canonical networks in the alpha and beta bands. Relative to broadband background brain activity, LSD significantly increases the prominence of high alpha networks (12.1 and 13.3 Hz) in all four experimental conditions, including resting-state and passive stimuli, and of high beta (25.3 Hz) in three conditions. However, LSD significantly decreased the prominence of low beta networks (18.1 and 19.3 Hz) in both the Open and Closed conditions and the prominence of low alpha (8.5 Hz) in just the latter.

Previous MEG studies on LSD and psilocybin found that both psychedelics decreased power in all sub-gamma frequency bands: delta, theta, alpha, and beta (Carhart-Harris et al., 2016; Muthukumaraswamy et al., 2013). This is consistent with a number of cross-species EEG studies that have shown that multiple psychedelics induce a broadband reduction in power, even at sub-hallucinatory doses in healthy organisms (Eidelberg et al., 1965; Horovitz et al., 1965; Hutten et al., 2024; Le et al., 2024; Murray et al., 2022; Rodin & Luby, 1966; Vejmola et al., 2021). That being said, it is worth noting that the psychedelics DMT and 5-methoxy-dimethyltryptamine (5-MeO-DMT) increase delta power in EEG recordings (Blackburne et al., 2024; Timmermann et al., 2023). LSD also diminishes evoked responses to light flash and to surprising audiovisual stimuli – that is, it decreases the mismatch negativity response – in both EEG and MEG (Chapman & Walter, 1965; Haggarty et al., 2024; Hutten et al., 2024; Murray et al., 2022; Timmermann et al., 2018).

Here, we measured the effect of LSD on the prominence of frequency-specific brain networks, not on their power. The FREQ-NESS technique in this study determines networks that maximally separate the covariance of individual frequencies from the broadband covariance, enabling us to quantify their prominence as the variance explained based on the associated eigenvalues. It is therefore well-suited for identifying the networks that are *uniquely* associated with specific frequencies. On the other hand, the power spectrum does not indicate the unique power at each frequency; it can be “contaminated” by contributions from broadband activity and/or noise. This broadband activity is typically reflected in the 1/f shape of power spectra, which determines the slope of the decay in power as frequency increases (Gerster et al., 2022; K. J. Miller et al., 2009). It is typically assumed that 1/f activity is aperiodic, arising from asynchronous neural potentials, so it is often labelled as noise (He, 2014; He et al., 2010). In order to isolate narrowband from broadband power, 1/f noise must be properly corrected for, ideally in a manner that accounts for the independence between noise and narrowband signal (Gyurkovics et al., 2021). As far as we are aware, there is only one EEG/MEG study on LSD that has corrected for 1/f noise (Muthukumaraswamy & Liley, 2018). Intriguingly, after correction, LSD appears to increase alpha power, highlighting the importance of removing broadband noise when determining power at individual frequencies. FREQ-NESS is inherently designed to isolate narrowband from broadband covariance, so it is ideal for measuring frequency-specific activity in situations where there is a substantial change in broadband noise, as is the case during the acute effects of psychedelics.

Furthermore, network prominence, as we defined it, is a fundamentally different quantity than power because it captures the extent to which multiple timeseries fluctuate together, rather than the amplitudes of individual timeseries. Our finding that LSD increases the explained prominence of high alpha and high beta is therefore compatible with, rather than contradictory to, previous results indicating that LSD reduces power in these bands. While LSD may lower the amplitudes of high-alpha and high-beta oscillations, it strengthens the connectivity within the networks formed by these oscillations. This may appear to be at odds with previous fMRI studies, which have found that psychedelics generally decrease within-network connectivity (Carhart-Harris et al., 2016; Dai et al., 2023; Madsen et al., 2021; Mason et al., 2020; Müller et al., 2018; Shinozuka, Jerotic, et al., 2024; Timmermann et al., 2023). However, unlike our methodology, fMRI analyses are unable to measure frequency-resolved connectivity due to their low temporal resolution and to the nature of the recorded signal. A prior MEG study used ICA to demonstrate that psilocybin significantly reduces within-network activation (Muthukumaraswamy et al., 2013). Other secondary analyses of the same MEG dataset in this study also found that LSD significantly reduced functional connectivity across all frequency bands, as measured by Granger causality and amplitude envelope correlations (Barnett et al., 2020; Pallavicini et al., 2019). These MEG studies averaged functional connectivity across whole frequency bands, rather than examining it at individual frequencies, as our method does. More importantly, ICA, Granger causality, and amplitude envelope correlations are affected by broadband power; if it decreases, then so do the amplitudes of the component timeseries. These techniques do not inherently account for the confounding influence of broadband noise, whereas FREQ-NESS does. Our study is the first to show that LSD’s effects on the strength of within-network connectivity depend on the frequency of oscillatory activity in these networks.

### Section 4.2. LSD disrupts the topographies of canonical alpha and beta networks

We also demonstrated that LSD substantially alters the spatial distributions or topographies of some of these networks. In particular, LSD shifts the low alpha (8.5 Hz) network in an anterior direction, away from the visual cortex and towards motor cortex. However, on LSD, the two high alpha networks (12.1 and 13.3 Hz) reverse direction – they become more localized to the visual cortex, especially at 12.1 Hz – except in the Video condition, in which the networks engage more of the sensorimotor cortex. The two low beta networks (18.1 and 19.3 Hz) encompass more of the temporal and occipital cortices on LSD, while the drug does not noticeably affect the topographies of the high beta networks (25.3 and 26.5 Hz).

Thus, in addition to altering the prominence of frequency-specific networks, which can be interpreted as resulting from their connectivity, LSD also reshapes their topographies. It is very well-established that alpha power (8-13 Hz) tends to be strongest in posterior brain regions, especially occipital cortex, where alpha plays a role in suppressing attention to and perception of visual input (Bazanova & Vernon, 2014; Berger, 1929; Klimesch, 1999). Higher alpha rhythms (12-14 Hz), also known as the “mu rhythm,” are localized to the sensorimotor cortex and inhibit motor responses (Roth et al., 1967; Sterman & Egner, 2006). In normal resting-state conditions, FREQ-NESS, both here and in previous applications, captures covariance in more posterior, occipital regions at lower alpha frequencies and covariance in more frontal, sensorimotor regions at higher alpha frequencies (Rosso et al., 2025). This may suggest that the mu rhythm accounts for more of the covariance at higher than lower alpha frequencies. However, we find that, on LSD, higher alpha frequencies are associated with greater occipital activation compared to placebo, though these frequencies still do engage sensorimotor areas. This shift, coupled with the anterior movement of lower alpha networks on LSD, indicates that the drug reverses the relationship between frequency and the topography of alpha networks. LSD appears to restructure the alpha networks such that lower frequencies exhibit more of the topography that is typically associated with the mu rhythm, whereas the topography of higher frequencies is more similar to that of the traditional, posterior alpha rhythm. (It is worth noting that, at both alpha frequencies, the networks are not neatly localized to either sensorimotor or occipital regions on LSD; each network contains a combination of both sets of regions.) The topography of lower alpha frequencies also encompasses temporal regions on LSD, but not at higher alpha frequencies, whereas the converse is true on placebo.

The functional significance of these topographical shifts is unclear and merits further investigation, especially given that the EEG/MEG literature in general tends to average activity across whole frequency bands rather than assessing individual frequencies. We propose two speculative explanations for our findings. First, the observed changes in brain topography may reflect a “flattening” of hierarchical processing under psychedelics. According to the RElaxed Beliefs Under pSychedelics (REBUS) theory, psychedelics reduce the influence of higher-order expectations (encoded as statistical priors) on brain activity. This enables sensory information to propagate from lower-to higher-order regions, along the cortical hierarchy, with less inhibition (Carhart-Harris & Friston, 2019). REBUS has received empirical support from other studies, which have confirmed not only that psychedelics flatten the hierarchy of the brain, but also that this effect predicts improvements in symptoms among patients with treatment-resistant depression (Deco et al., 2024; Girn et al., 2022; Mediano et al., 2024; Schartner et al., 2017; Shinozuka et al., 2025; Timmermann et al., 2023). REBUS proposes that reductions in alpha power on psychedelics may reflect a flattened hierarchy of information flow in the brain because alpha generally encodes top-down inhibition of sensory representations (Carhart-Harris & Friston, 2019; Jensen, 2024; Klimesch et al., 2007; E. K. Miller et al., 2018). Our study illustrates that LSD alters the canonical topography of alpha networks, which may disrupt the ability of alpha oscillations to inhibit sensory cortices in a top-down manner. In particular, at high alpha frequencies, the visual areas that are inhibited by alpha are more posterior on LSD than placebo. This dysregulation of top-down alpha activity may diminish hierarchical information processing. However, while compelling, this relationship between the flattening of brain hierarchies and the reshaping of alpha networks is speculative and should be directly investigated in future research.

Secondly, the observed shifts in topography may arise from disruptions in thalamocortical connectivity. Posterior alpha oscillations are generated by the pulvinar and/or lateral geniculate nucleus, two nuclei of the thalamus that fire at the alpha rhythm (Fiebelkorn et al., 2019; Hughes & Crunelli, 2005; Lőrincz et al., 2009; Saalmann et al., 2012). Mu alpha oscillations in sensorimotor cortex are produced by ventral posteromedial and posterolateral nuclei in the thalamus, relaying signals from the brainstem (Jones et al., 2009). In rodents, LSD reduces the spiking of GABAergic neurons in the thalamic reticular nucleus (TRN), a sheet that surrounds and “gatekeeps” the thalamus, leading to disinhibition of the thalamus (Inserra et al., 2021; Onofrj et al., 2023). Hence, both fMRI and MEG studies in humans have demonstrated that psychedelics dramatically elevate functional connectivity between the thalamus and the cortex, especially enhancing the connectivity of the pulvinar and ventrolateral nuclei (Avram et al., 2022; Bedford et al., 2023; Coleman et al., 2024; Delli Pizzi et al., 2023; Müller et al., 2017; Preller et al., 2018; Tagliazucchi et al., 2016). Thus, LSD may perturb the thalamic inputs that sustain normal alpha and mu rhythms, which could alter the topography of alpha networks in a frequency-dependent manner. That is, LSD could interfere with the posterior distribution of alpha and mu activation at particular frequencies, as we discovered at both 8.5 Hz and 12.1/13.3 Hz.

Furthermore, these perturbations may impair the ability of the thalamus to perform its role of gating sensory information to the cortex and filtering out unimportant stimuli (McCormick & Bal, 1994; Vollenweider & Geyer, 2001). Thalamocortical connectivity may therefore underlie the subjective experience of “aberrant salience” on psychedelics: everything, including things that would normally seem mundane, feels profoundly significant (Hendricks, 2018; Stace & Smith, 1987). Hence, increases in thalamocortical connectivity are correlated with altered states of consciousness experienced on psychedelics, including feelings of “oceanic boundlessness” or vast interconnectedness (Müller et al., 2017). The thalamocortically mediated disruption of posterior alpha at either 8.5 Hz (decreased alpha prominence) or 12.1/13.3 Hz (increased alpha prominence) could also play a role in the visual hallucinations that occur on psychedelics. Disinhibition of the TRN and structural alterations of the pulvinar are thought to cause visual hallucinations in dementia with Lewy bodies (DLB), which could be related to alterations in posterior alpha in this condition (Babiloni et al., 2018; Delli Pizzi et al., 2023; Esmaeeli et al., 2019; Rosenblum et al., 2020). (That being said, DLB visual hallucinations are phenomenologically distinct from those of psychedelics (Esmaeeli et al., 2019).) However, FREQ-NESS does not establish a causal relationship between thalamocortical connectivity and the topography of low or high alpha networks, so our interpretation remains speculative.

A unique feature of the dataset in this study is the acquisition of MEG recordings during both resting-state and passive stimulus conditions. We noticed that the topographies in the Video condition tended to diverge from those of the other conditions. In particular, whereas LSD shifted the topographies of high alpha networks in a posterior direction in the Open, Closed, and Music conditions, the topographies moved in an anterior direction in the Video condition at both 12.1 and 13.3 Hz. In the Video condition on LSD, the networks at these frequencies recruited the primary motor and somatomotor cortex, as well as other frontal areas, while disengaging visual cortex. Since alpha generally suppresses visual perception, lack of alpha covariance in occipital areas in the Video condition may suggest that LSD heightens visual alertness, but only when visual stimuli are being presented and not during resting-state or auditory stimuli.

In the Open and Closed conditions, LSD shifts the topography of not only alpha but also low beta prominence. While some have proposed that the beta rhythm is generated in cortex through intrinsic membrane time constants, such as the kinetics of the M-type potassium current in excitatory neurons, or subcortical structures like the basal ganglia (Bevan et al., 2002; Holgado et al., 2010; Jones et al., 2009; Kopell et al., 2000; Leventhal et al., 2012; McCarthy et al., 2011; Pinto et al., 2003; Roopun et al., 2006), a recent model of MEG data implicates the thalamus as one of the drivers of the beta rhythm (Sherman et al., 2016). In this model, spontaneous beta activity in the sensorimotor cortex arises from two nearly synchronous events: weak and strong excitation of proximal and distal dendrites of pyramidal neurons, respectively, with the latter lasting the duration of one beta cycle. Projections from the thalamus are hypothesized to provide the excitatory drive to the distal pyramidal dendrites, thereby facilitating the beta rhythm. By enhancing the transmission of neural signals along these thalamocortical projections, LSD may augment the excitatory drive that helps to generate the beta rhythm, potentially enhancing beta in brain regions where it typically is not produced. In particular, because LSD elevates the functional connectivity between the pulvinar and cortex, and certain segments of the pulvinar are strongly coupled with temporal and occipital cortices (Arcaro et al., 2018; Delli Pizzi et al., 2023), LSD may increase beta prominence in temporo-occipital regions. This is consistent with the topographies of low beta networks that we observed in this study, which expanded on LSD into left superior and middle temporal gyri, as well as the left middle occipital gyrus, at multiple frequencies and conditions. Intriguingly, aberrant functional connectivity and structural morphology of the left superior temporal gyrus may underlie auditory hallucinations in conditions like schizophrenia (Barta et al., 1990; Hugdahl et al., 2009). While hallucinations on LSD tend to be more visual than auditory, both psychedelic-induced and schizophrenic hallucinations share the common characteristic of misattributing internally generated perception to the external world (Hugdahl et al., 2009). For instance, in both conditions, people may feel as though spiritual beings are communicating something of profound significance to them (Griffiths et al., 2019; Grover et al., 2014).

### Section 4.3. Limitations

The eigenspectrum – the plot of frequency vs the leading eigenvalue – exhibits a clear 1/f slope in all conditions except Closed. This is likely attributable to the fact that alpha oscillations become much more prominent whenever eyes are closed (Berger, 1929), leading to a stronger alpha peak both in the traditional power spectrum and the eigenspectrum here. It is surprising that the eigenspectrum has a 1/f slope in the Music condition, where participants’ eyes are also closed. We initially thought that this may be related to the very slow rhythm of the music, which was excerpted from the ambient album *Yearning* by Robert Rich and Lisa Moskow; lower frequencies in the music are much more common than higher frequencies. Previous applications of FREQ-NESS found that the variance explained by low-frequency, 2.4 Hz activity in the brain significantly increases relative to resting-state when a 2.4 Hz auditory stimulus is played (Rosso et al., 2025). However, none of the peaks in the power spectrum of the music in this study corresponded to the frequencies that exhibited significant between-drug differences in leading eigenvalue. Future research should consider playing simple, single-tone auditory stimuli to participants during the acute effects of psychedelics, rather than ambient music that lacks a clear beat, in order to determine whether LSD alters the topography of brain networks at the frequency of the stimulus.

Brain activity in the LSD Music condition may have been noisier than the other LSD conditions, as the number of epochs that were removed due to artefacts was significantly higher compared to placebo in this condition. However, there was no significant between-drug difference for the other conditions. Music was played while participants’ eyes were closed, while the video was presented while participants’ eyes were open, which complicates the comparison between these two conditions.

## Section 5. Conclusion

In the current study, we measured the acute effects of LSD on frequency-specific brain networks with a simple yet powerful technique, FREQ-NESS, which is capable of separating covariance at individual frequencies from broadband covariance. We report two major findings. Firstly, LSD significantly increased the prominence of high alpha (12.1 and 13.3 Hz) and high beta (25.3 Hz) networks in both resting-state and passive stimulus conditions, relative to broadband covariance. While LSD is known to reduce alpha power, the enhanced prominence of high alpha networks indicates that the drug may actually strengthen the connectivity of networks oscillating at 12.1 and 13.3 Hz. LSD also significantly decreases the prominence of low alpha (8.5 Hz) and low beta (18.1 and 19.3 Hz) networks in some conditions. Secondly, LSD reshaped canonical resting-state networks in a frequency-dependent manner. The drug shifted alpha networks, which are typically occipital, in an anterior direction at low alpha frequencies and in a posterior direction at some high alpha frequencies. On LSD, low beta networks, which are usually localized to sensorimotor cortex, engage more temporo-occipital brain regions. We encourage future researchers to link the hallucinogenic and therapeutic effects of psychedelics to network connectivity at specific frequencies, using carefully controlled experimental designs. Moreover, building on the substantial evidence of interactions between brain regions in different frequency bands, such as cross-frequency coupling and other forms of coordination (Canolty & Knight, 2010; Jensen & Colgin, 2007) future research should further explore the interactions between the frequency-specific whole-brain networks identified in the present study. This would provide deeper insights into how LSD reshapes the dynamic interplay between brain networks.

## Notes

### Competing Interest Statement

The authors have declared no competing interest.

