## Supplementary Materials for "LSD reconfigures the frequency-specific network landscape of the human brain"

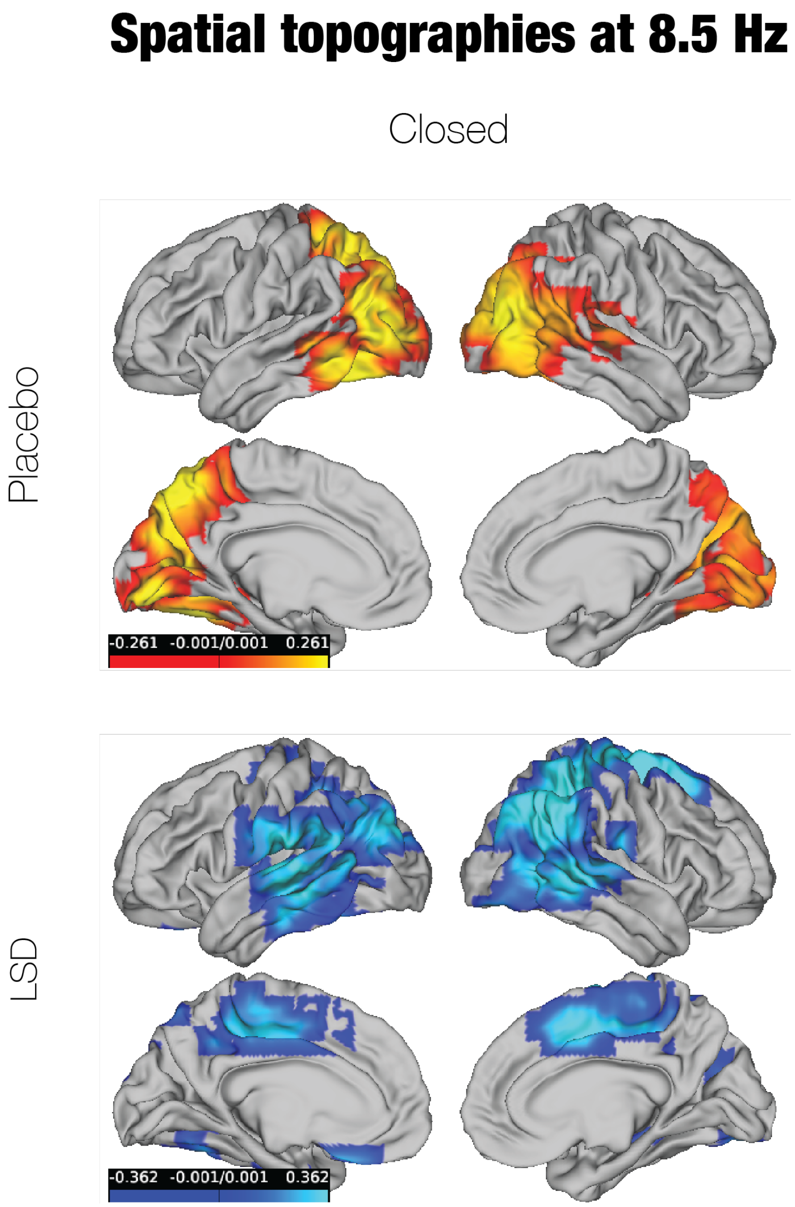


***Figure S1. In the Closed condition, LSD engages more anterior regions in the network associated with 8.5 Hz activity.*** *These include the right primary motor cortex, right primary somatosensory cortex, right supplementary motor area, bilateral middle cingulate, and various bilateral temporal cortices. Meanwhile, the network becomes less active in visual areas like bilateral V1, bilateral V2, and right superior and middle occipital gyrus. The Closed condition was the only condition in which LSD caused a significant change in the amount of variance explained by the leading component.*

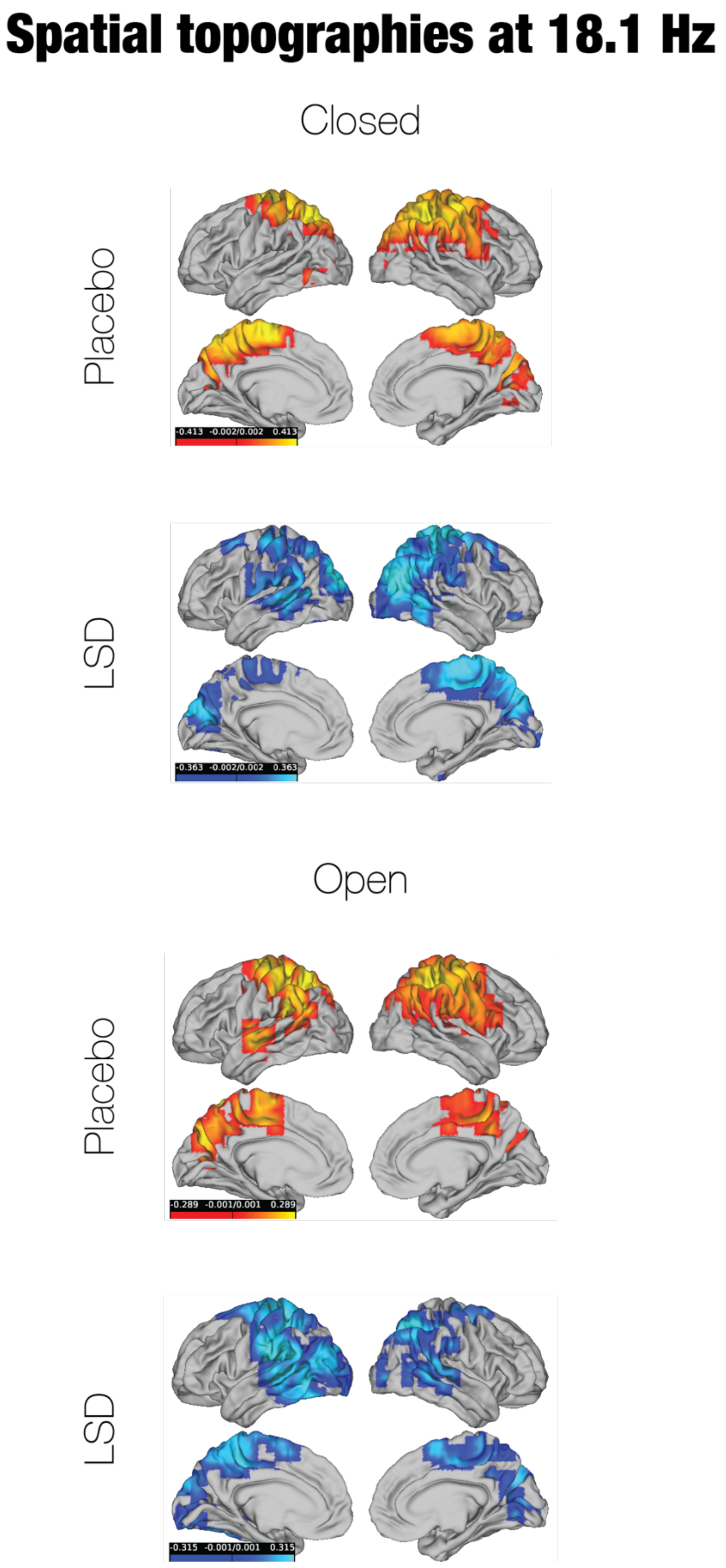


***Figure S2. LSD shifts the network associated with 18.1 Hz activity towards the temporal and occipital lobes in the non-stimulus conditions.*** *At this frequency, LSD significantly decreases the amount of variance explained by the leading component in only the Closed and Open conditions. LSD recruits temporo-occipital regions, including the bilateral middle temporal gyrus and left middle occipital gyrus, into the leading component at 18.1 Hz. In the right hemisphere, the network retreats from frontoparietal areas, including the primary motor cortex and somatosensory cortex and inferior parietal gyrus.*

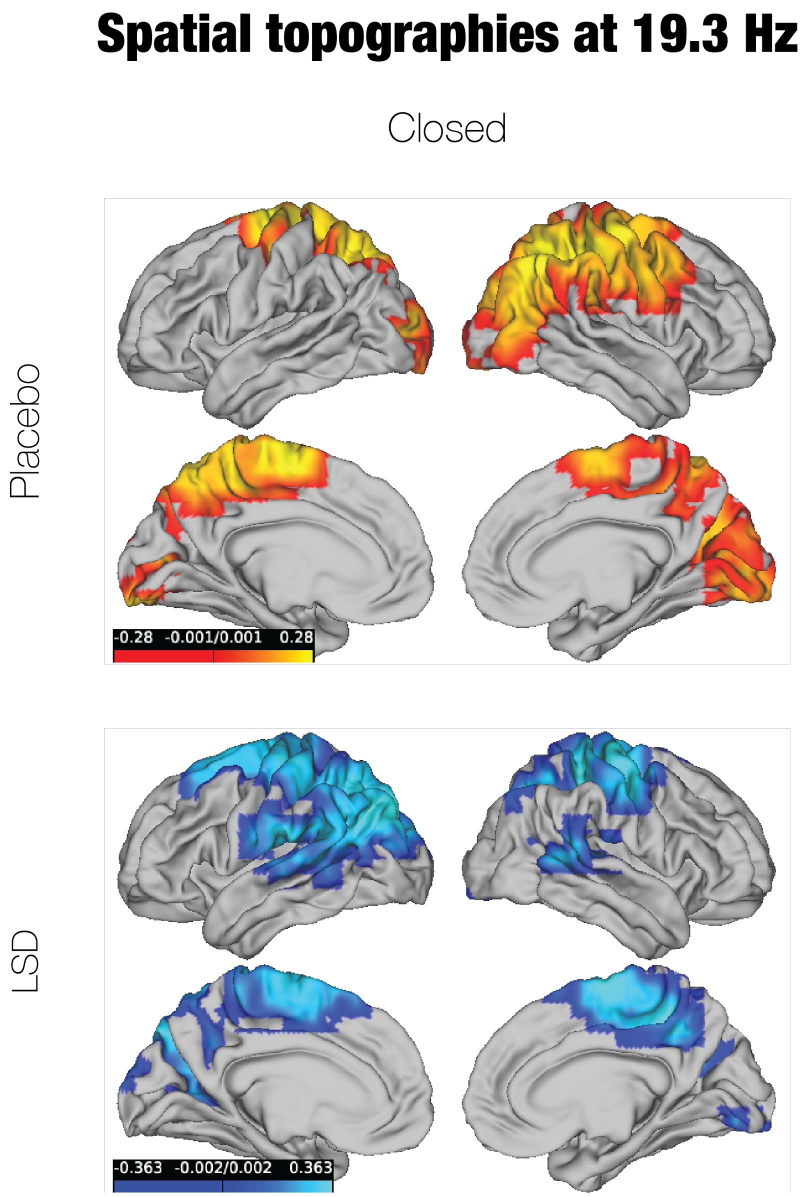


***Figure S3. In the Closed condition, LSD again engages temporal and occipital regions in the network associated with 19.3 Hz activity.*** *LSD significantly reduces the variance explained at this frequency in only the Closed condition. The leading component expands into the left superior and middle occipital gyri, though it withdraws from the same visual areas in the right hemisphere, and the left superior and middle temporal gyri. Additionally, the network encompasses the parietal lobe, left primary motor and somatosensory cortex, and bilateral supplementary motor area,*

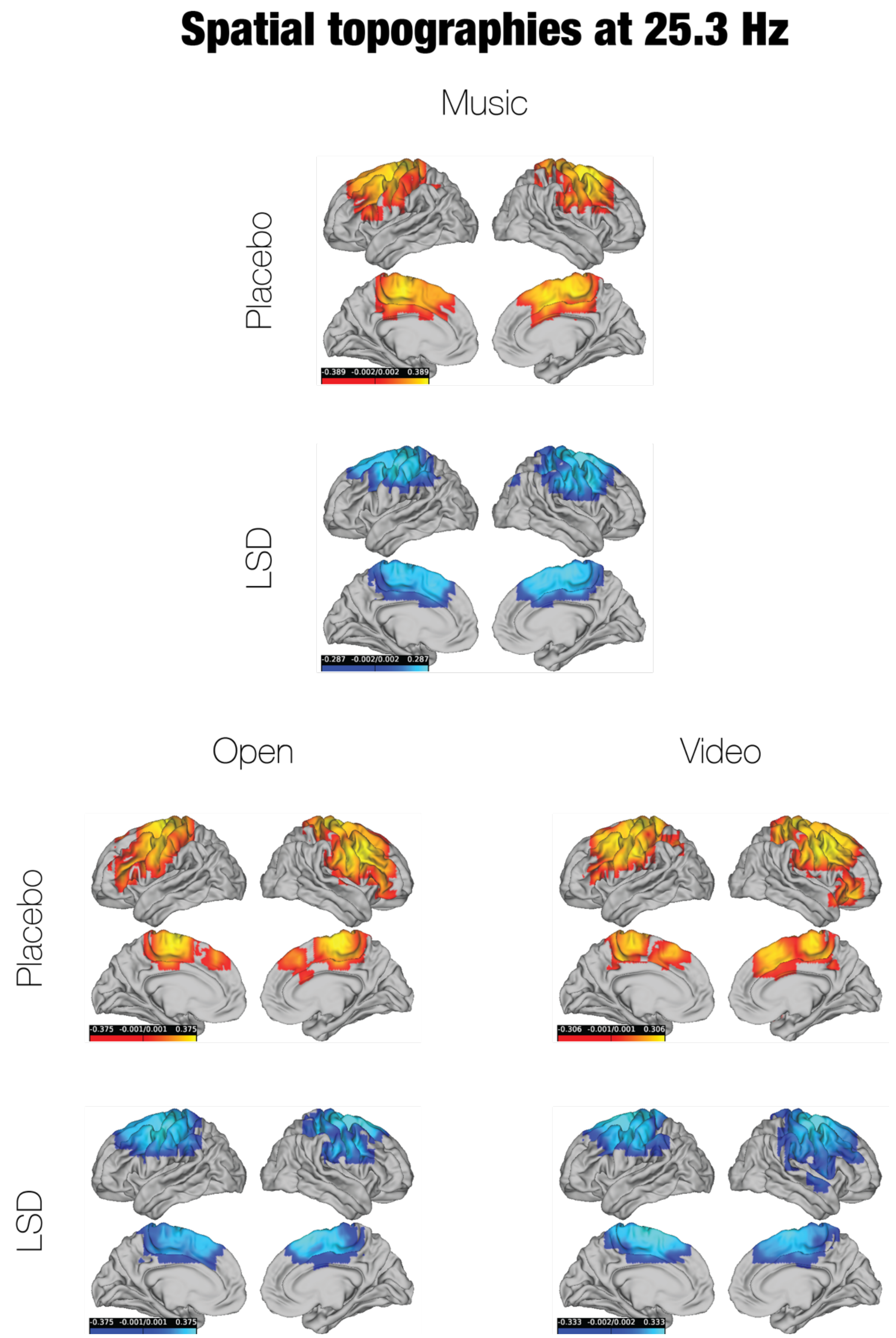


***Figure S4. In the Open, Music, and Video conditions, LSD does not have a strong effect on the spatial topographies of the networks associated with 25.3 Hz activity.*** *LSD significantly increased the variance explained by 25.3 Hz in all conditions except Closed. However, across the three conditions, the corresponding networks look very similar between placebo and LSD. The networks are concentrated in motor and somatosensory cortex and other frontal areas.*

### Spatial topographies at 26.5 Hz

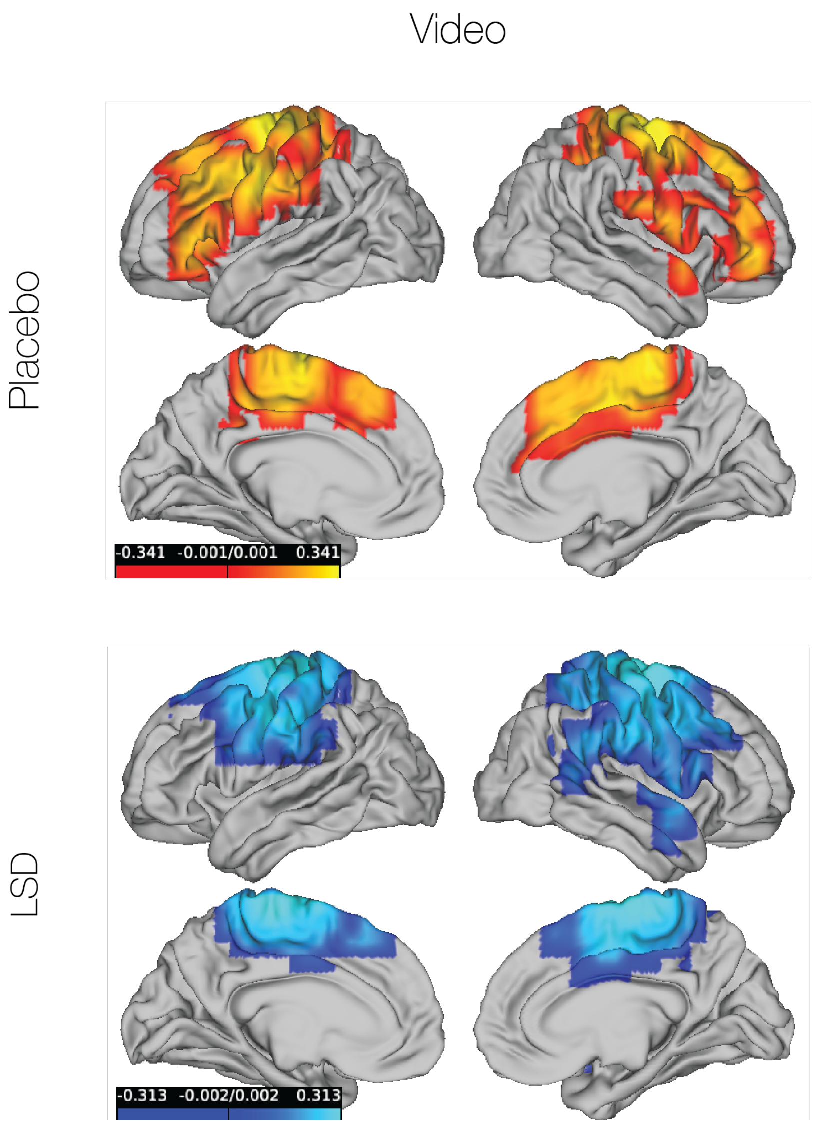


***Figure S5. LSD moves the 26.5 Hz network away from some frontal regions in the Video condition, but otherwise does not substantially alter the topography of the network.*** *These include the bilateral superior and middle frontal gyrus, left inferior frontal gyrus pars triangularis, and left primary motor cortex. The network does become more activate in the bilateral primary somatosensory cortex. The Video condition was the only condition in which LSD significantly altered the variance explained by the leading component at 26.5 Hz.*
